# Reported estimates of human airway dimensions are inconsistent across studies

**DOI:** 10.64898/2026.01.19.699966

**Authors:** Mahesh K. Ola, Morgan Seal, Raza A. Sarwar, Sai K. Mattireddy, Chris E. Brightling, Kelly Burrowes, Himanshu Kaul

## Abstract

**Rationale:** Respiratory diseases are a source of immense socioeconomic burden globally. *In silico* approaches can predict changes in human lung function due to disease or response to therapy. By stratifying patient-specific response *a priori*, these models can enable clinical-scale deployment of precision medicine strategies. Key to this is developing accurate organ geometries on which the models can be simulated. However, we lack analyses assessing the clinical applicability of reported airway dimension estimates.

**Objective:** To investigate physiologically-/anatomically-relevant airway dimension estimates and evaluate consistency across reported literature.

**Methods:** We conducted a systematic review of 37 published datasets. Airway wall thickness estimates were mined for healthy subjects and patients, and standardised to the Horsfield order airway generations. We simulated dynamic lung function to quantitatively assess their physiological relevance. We created an online database to make all datasets available to the research community.

**Measurements and Main Results:** Reported human airway wall thickness estimates are inconsistent across studies. K-means clustering divided estimates for healthy subjects and patients into three and four clusters, respectively. Only one of the clusters in each category yielded anatomically-relevant estimates. Pressure-volume curves generated to assess physiological relevance also showed that only one cluster in each category exhibited plausible physiology. Principal Component Analysis weakly implicated imaging modalities to explain this inconsistency.

**Conclusions:** Reported airway dimension estimates are inconsistent and lack standardisation. To support future modelling efforts, we report physiologically-relevant estimates and introduce an open-access airway-dimension database to help standardise geometric inputs and quantify how measurement variability propagates to functional predictions.

## INTRODUCTION

Asthma affects ∼300 million people worldwide [1, 2] with a high mortality rate (∼1000 deaths/day). Asthma is characterised by heterogeneity in clinical phenotype, severity, pathobiology, and response to therapy. Consequently, existing therapeutic methods fail to effectively manage asthma [3]. Precision medicine, which entails tailoring therapies to patients’ unique profiles, is a potentially powerful approach to treating asthma. However, this requires 1) precise understanding of the multiscale mechanisms that mediate multiple asthma phenotypes and aetiologies 2) mapped on to patient-specific airways with high spatiotemporal accuracy.

There is a significant corpus of work, including by us, done along both strands. For example, we have previously developed an agent-based [4] virtual asthma patient [5, 6] that captured asthma pathogenesis based on interactions between epithelial, mesenchymal, and inflammatory agents (or virtual cells). In addition to capturing the mechanisms that mediate eosinophilic asthma, the model accurately captured response to fevipiprant [6], mepolizumab [6], and benralizumab [5]. Several groups have developed patient-specific computational models at the macro-scale, simulating various aspects of lung function, including ventilation [7–12], particle transport [13–15], perfusion [16], gas exchange [17], and more for healthy subjects and diseases lungs. These types of models are patient-specific, or patient-based, in terms of their anatomy or the geometry used to represent the appropriate pulmonary structures. The two varieties of models upon integration can help couple (spatiotemporally) cellular interactions with tissue/organ mechanics/function and, potentially, capture asthma pathobiology and stratify response to therapy at a patient-specific level [18].

To develop these in silico models, the starting point entails creating the virtual geometry of the lung airway. This is typically achieved by imaging the lung (via computed tomography (CT), high-resolution CT (HRCT), multi-detector CT (MDCT), Magnetic Resonance Imaging etc), followed by segmentation and 3D reconstruction (including noise reduction, mesh smoothening, etc), and construction of a computational mesh that can be used in simulations. The underlying biophysical equations can then be solved, using mathematical representations and computational software, to determine the relevant lung function. Swan et al. (2012) [7] presented a model of ventilation within a subject-based anatomical geometry representing a healthy human lung. In addition to using image derived airway geometry, a volume-filling branching algorithm [19] can be used to ‘grow’ the distal airways, which are unable to be resolved from imaging. This flow (ventilation) model represents the full conducting airway tree with each acinus represented as a lumped compliant unit, driving airflow into the model. Key to virtual lung geometry construction is precise determination of airway dimensions (e.g. lumen diameter and wall thickness) achieved via centreline extraction and analysis, including model-based fitting (as needed). These image-based virtual airway geometries, and estimated airway dimensions, serve as the foundation for *in silico* models that compute mechanical and functional changes vs ‘baseline’ due to disease/therapy. Estimates of airway dimensions are, therefore, critical to the models’ prediction accuracy. Yet, there is a lack of systematic analysis on existing estimates of airway dimensions and how precisely they reflect the general anatomy of a (healthy/asthmatic) individual.

Here, we asked: is there a standardised estimate of (healthy/diseased) airway dimensions that can be used to generate a virtual airway tree to simulate clinically-relevant changes in lung mechanics and function? We addressed this question via a meta-analysis of published airway dimension estimates, which included standardising all datasets, conducting unsupervised clustering, and simulating lung function for each cluster (where possible). Our work reveals physiologically-relevant wall thickness estimates (for health and disease) that modellers can use to simulate lung mechanics and function. To ensure broad accessibility, we created an open-access database, The Human Airway Registry, where researchers can upload their estimates to visualise/compare against existing studies, determine whether their choice of estimates are physiologically-relevant, or download physiologically-relevant estimates to create virtual geometries for their models.

## RESULTS

### Reported dimension estimates are highly inconsistent for both healthy subjects and patients

We gathered and reviewed 45 datasets in total (22 healthy subjects (*H*); 15 with respiratory diseases (*D*), specifically asthma and COPD; and eight studies where the disease status was not clearly identified (*U*)) published over four decades (Tables 1). Given lack of clarity around disease status, we did not consider the *U* datasets in our analysis. To enable direct comparison of consistency, we next standardised dimensions across the datasets by normalising wall thickness estimates to the Horsfield order (see Methods). We found the estimates to be highly inconsistent. Figures 1A-D show the standardised estimated wall thickness and coefficient of variation (CoV) for each airway generation (health and disease). The absolute estimates vary by at least an order of magnitude and display very high CoV across all datasets. As expected, the airways/generation were thicker compared to the healthy cases (Figure 1E).

**Figure 1:**
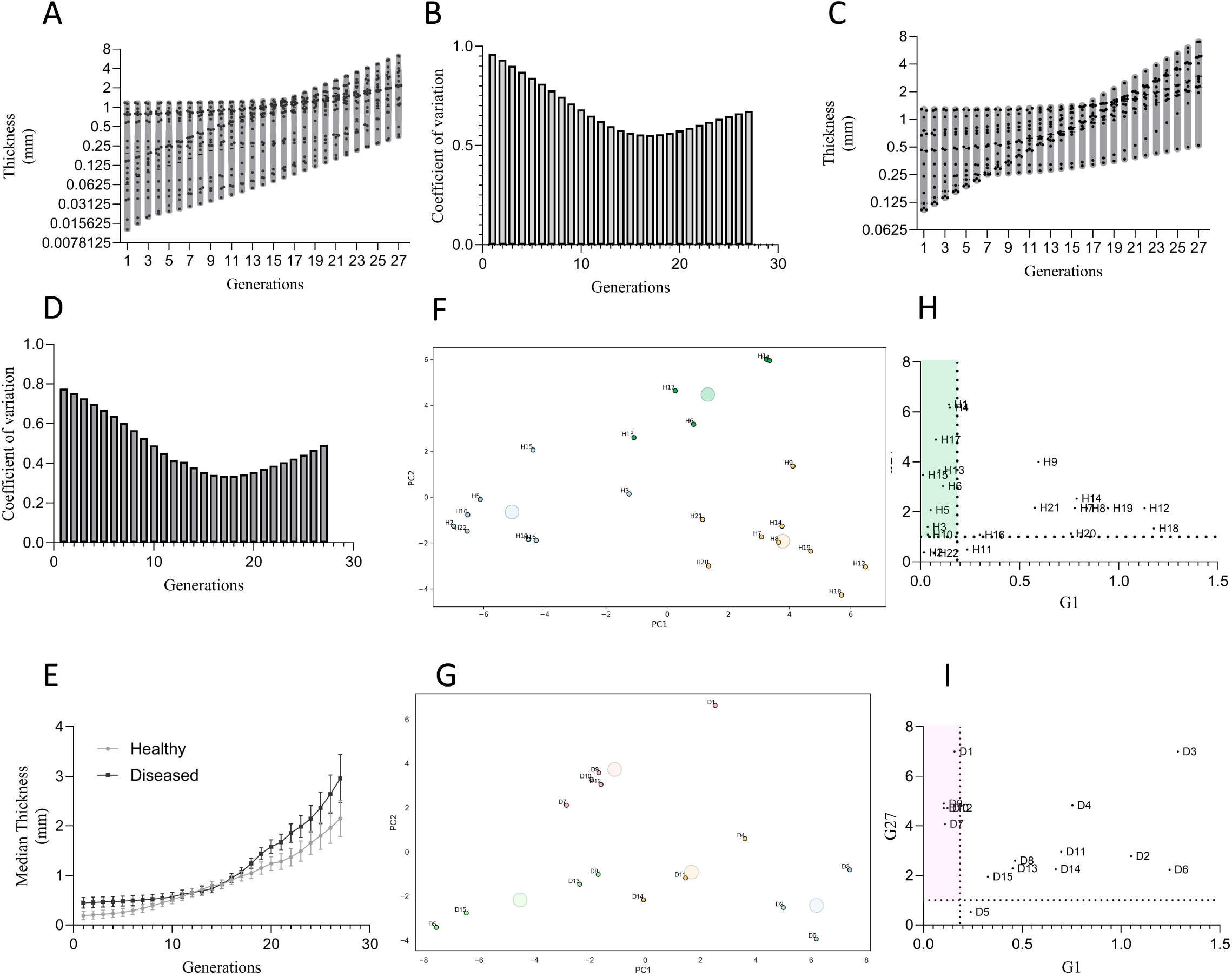
**(A)** Violin plot shows estimated airway wall thickness for all airways standardised to Horsfield order for the healthy datasets. Each dot represents a dataset. **(B)** Histograms show the coefficient of variation (CoV) for wall thickness estimates across all Horsfield airway generations for the healthy datasets. **(C)** Violin plot shows estimated airway wall thickness for all airways standardised to Horsfield order for the diseased datasets. Each dot represents a dataset. **(D)** Histogram shows the coefficient of variation (CoV) for wall thickness estimates across all Horsfield airway generations for the diseased datasets. **(E)** Median wall thickness for healthy vs diseased airways. Bars represent standard error. **(F)** PCA projection with K-means clustering of wall thickness estimates for all 27 airway generations for healthy datasets. **(G)** PCA projection with K-means clustering of wall thickness estimates for all 27 airway generations for diseased datasets. **(H)** G1 wall thickness vs G27 wall thickness for healthy datasets. The shaded region (green) shows the anatomically-relevant datasets, which form the green cluster in the PCA projection for healthy datasets. **(I)** G1 wall thickness vs G27 wall thickness for diseased datasets. The shaded region (pink) shows the anatomically-relevant datasets, which form the pink cluster in the PCA projection for diseased datasets.

To identify datasets with similar wall thickness estimates, we conducted PCA on the estimated wall thickness followed by K-means clustering (health and disease, separately, Figures 1F,G). The elbow method suggested data to be spread across three clusters for healthy subjects and 4 clusters for patients. We next asked which of these clusters are anatomically-relevant. To achieve this, we plotted airway wall thickness in generation (G) 1 vs wall thickness in G27 for each dataset (Figures 1H,I). Datasets either yielded estimates that were either unpractically low for G27 (bottom left corner of the plot) or high for G1 (right half of the plot). The estimates shaded green (Figure 1H) showed anatomically relevant estimates for health, which belonged to Cluster II (green) on the PCA plot (Figure 1F). For patient datasets, the estimates shaded pink (Figure 1I) showed anatomically relevant estimates, which belonged to Cluster II (pink) on the PCA plot (Figure 1G). The anatomically-relevant clusters were localised ∼0 on PC1 and >2 on PC2 for both health and patient datasets.

### Variance in wall thickness estimates is (weakly) explained by the choice of imaging modality

We next investigated the reason for this unusually high variance and inconsistency in wall thickness estimates. Specifically, we considered factors such as mean age of subjects/patients, sex differences (whether studies were male or female dominated or equally represented), publication year of the study, and the imaging modality used. Table 1 details these metadata for each dataset. We next plotted the airway wall thickness estimates and associated CoV for each of those attributes to assess whether the anatomically-relevant (health or disease) cluster were enriched for a specific attribute.

**Table 1:**
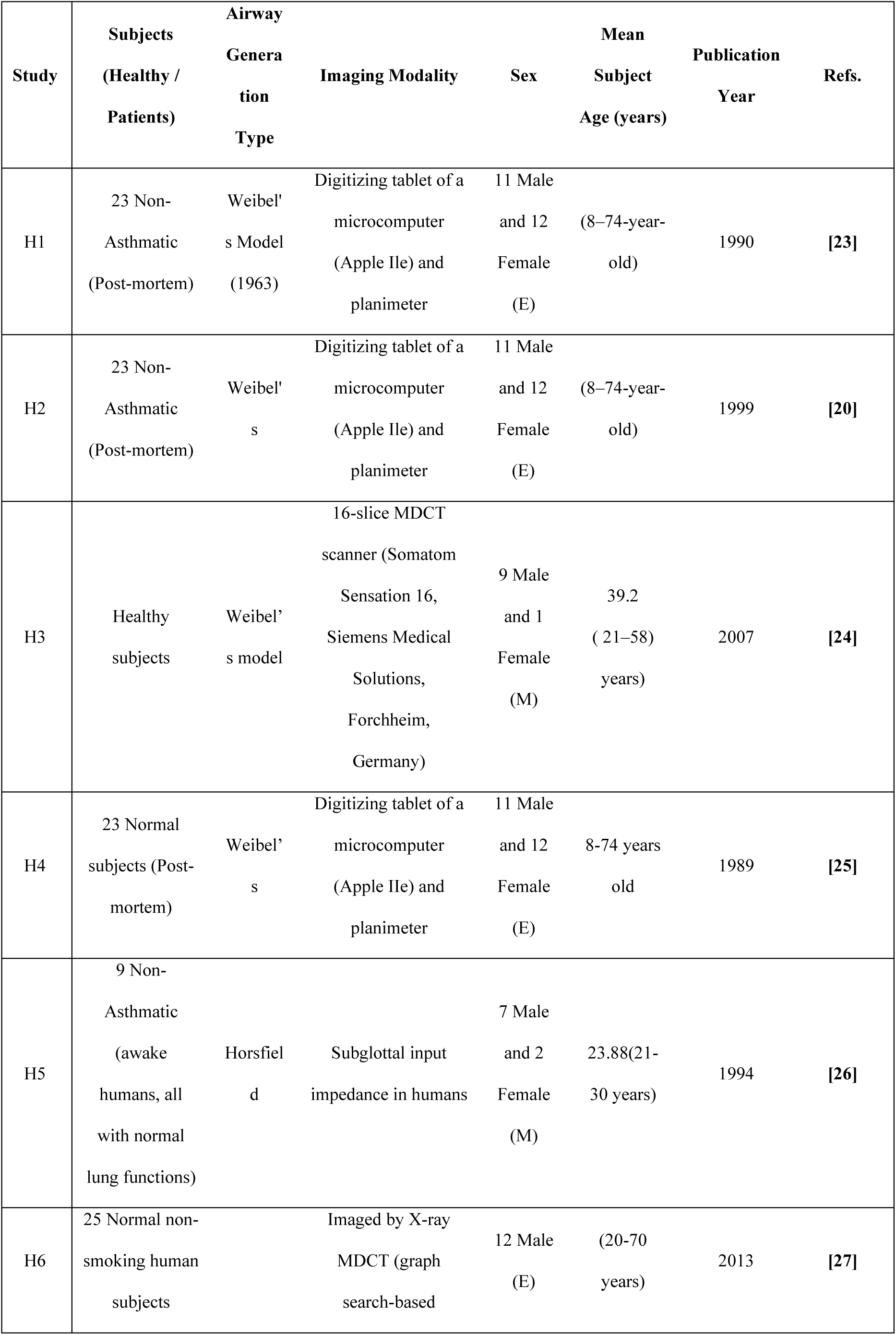
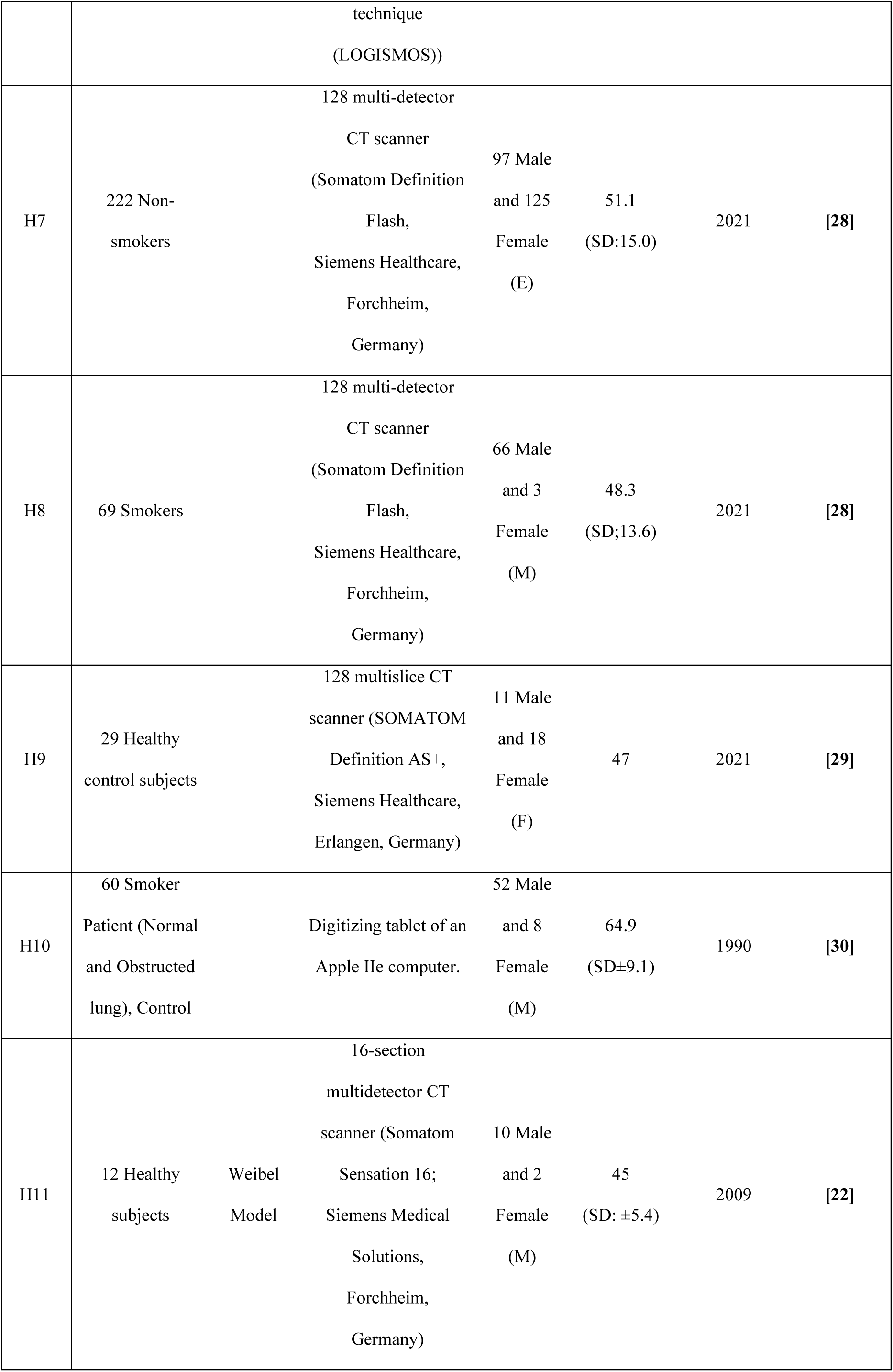
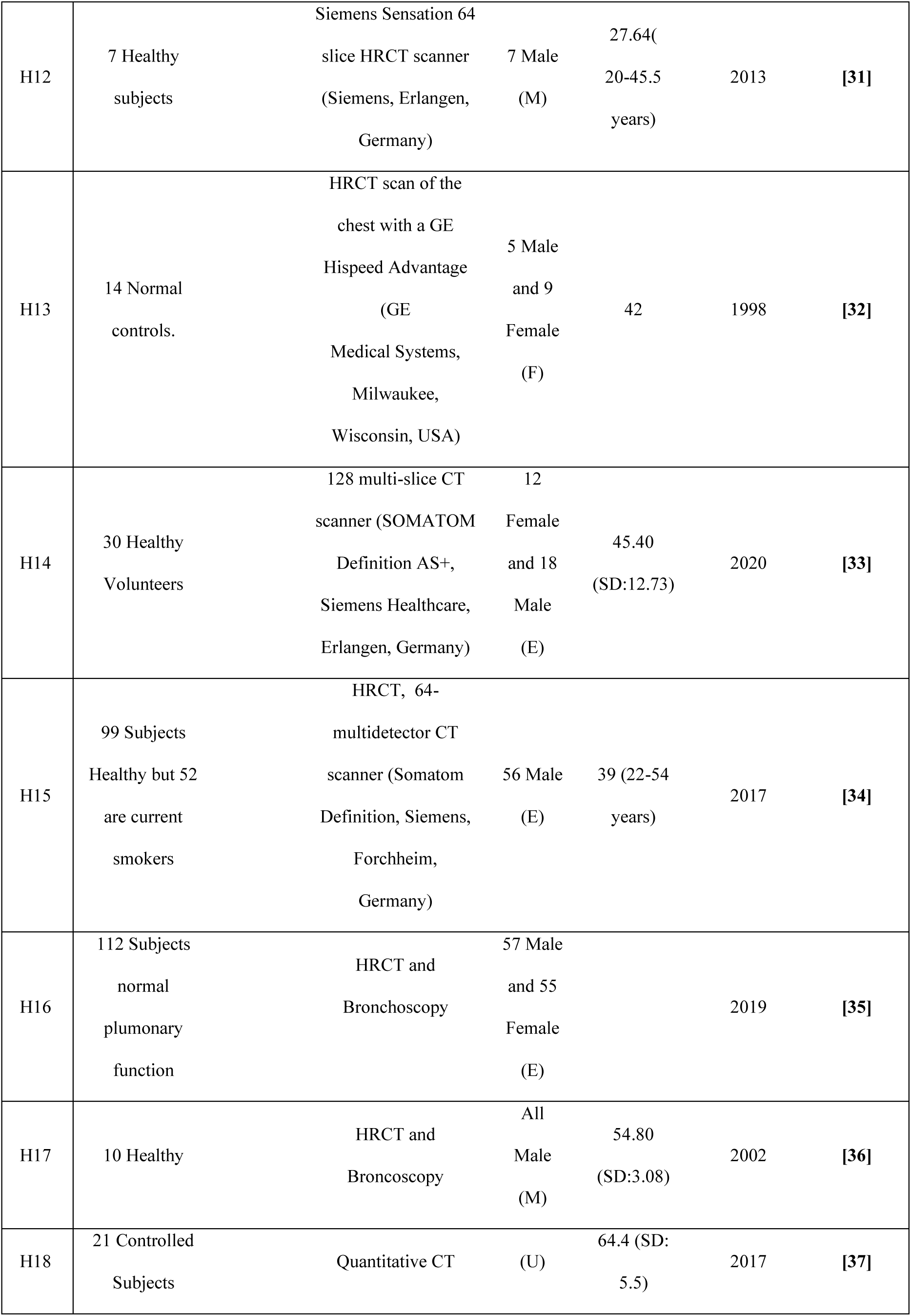
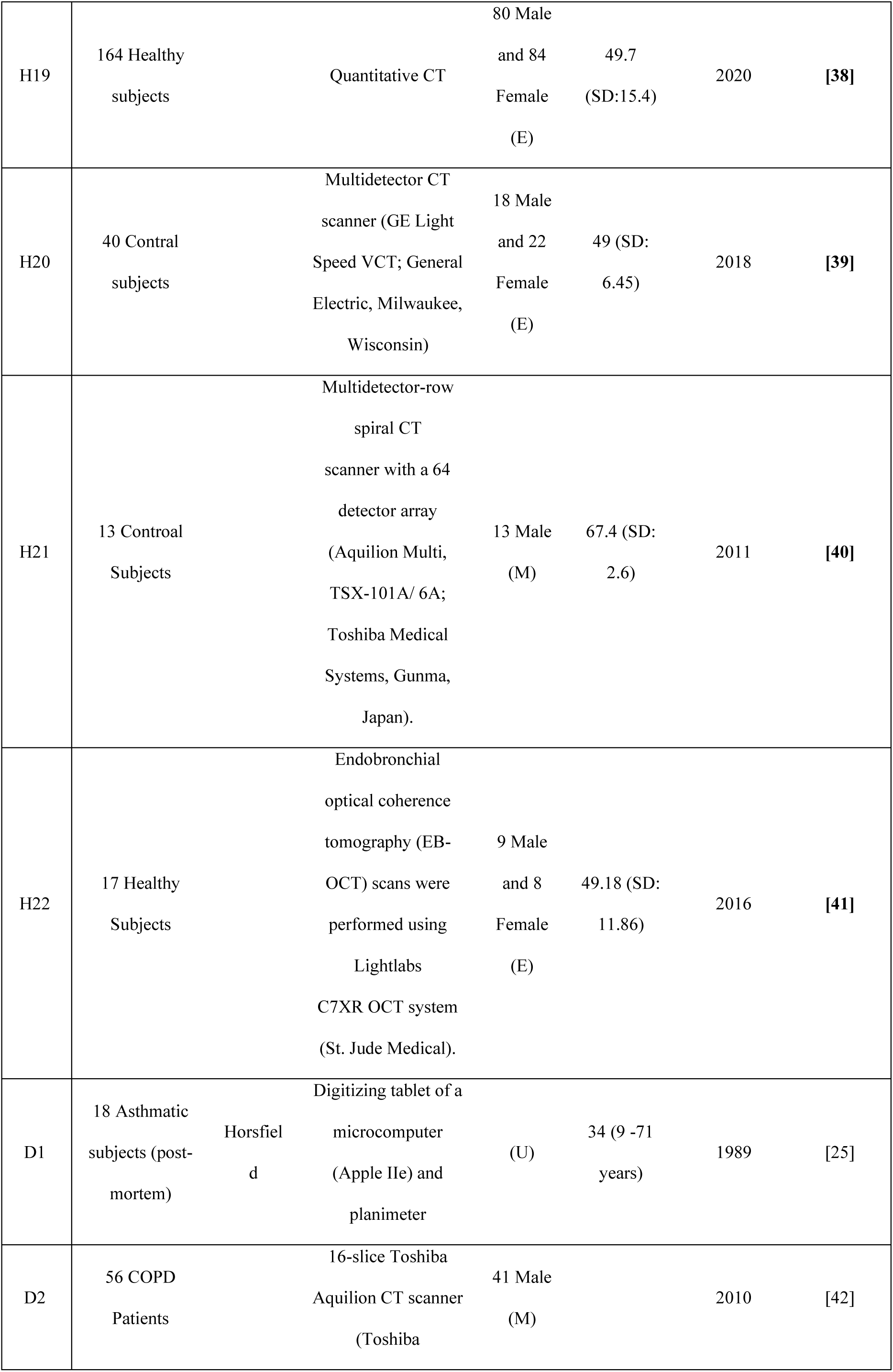
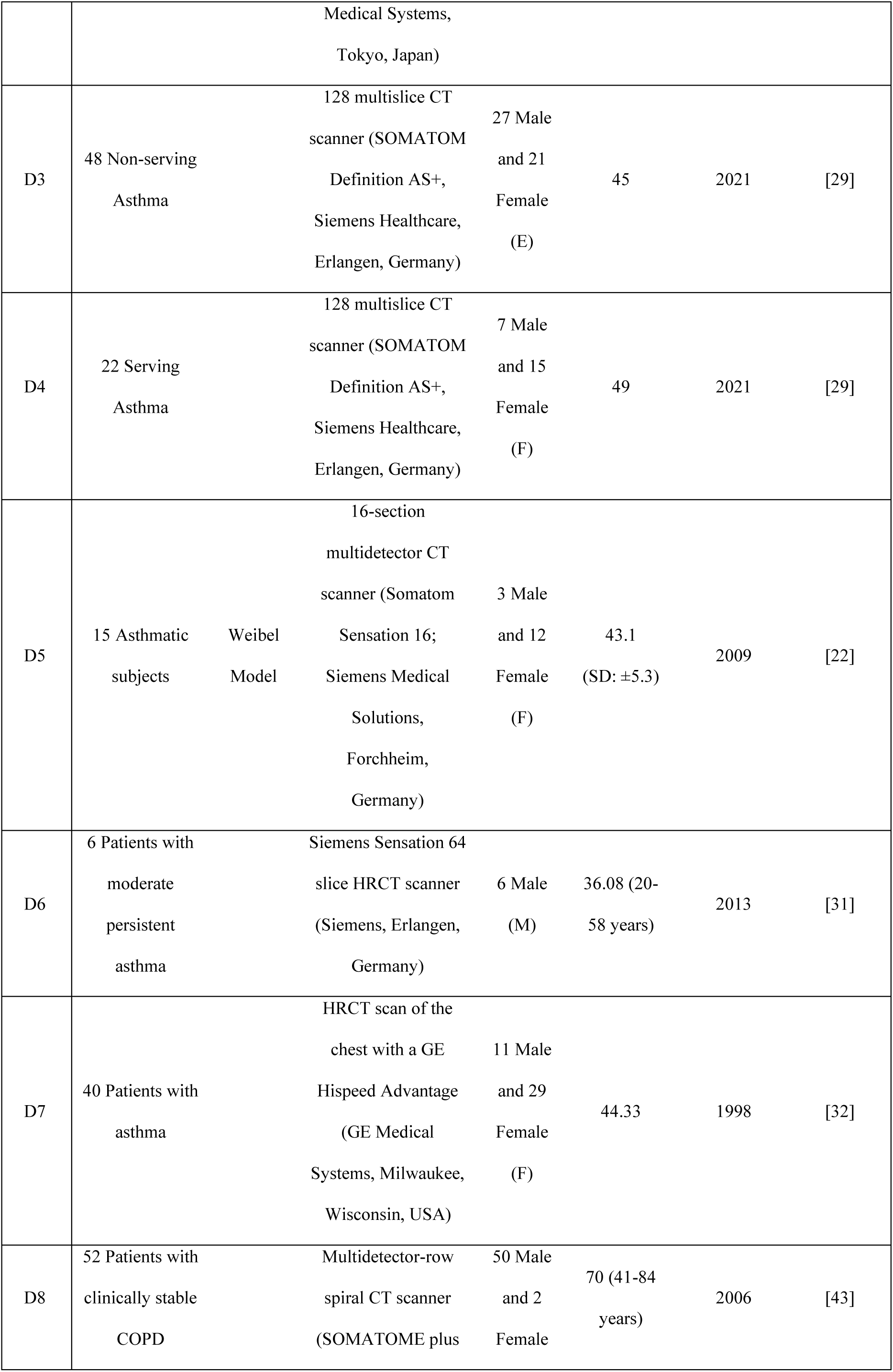
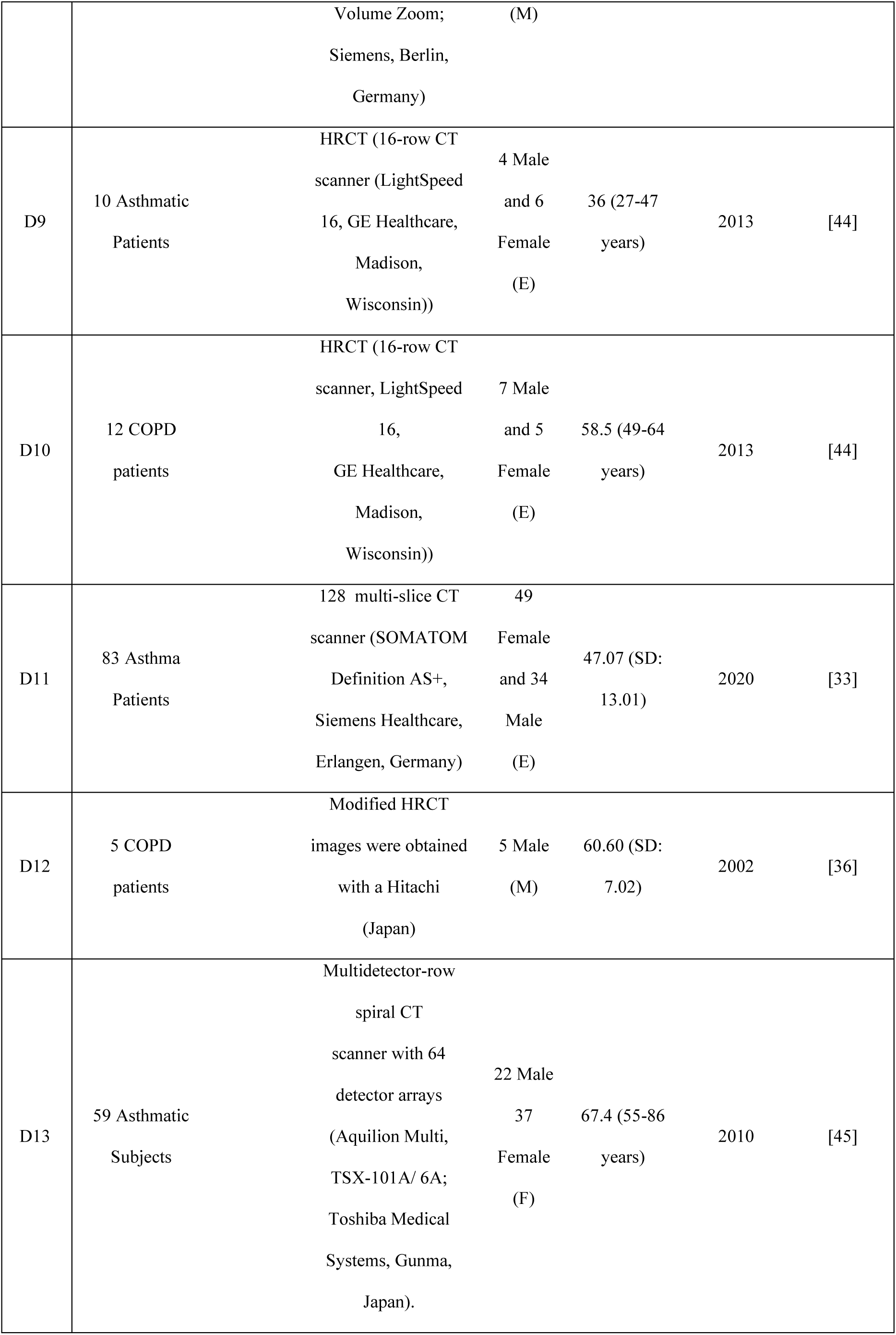
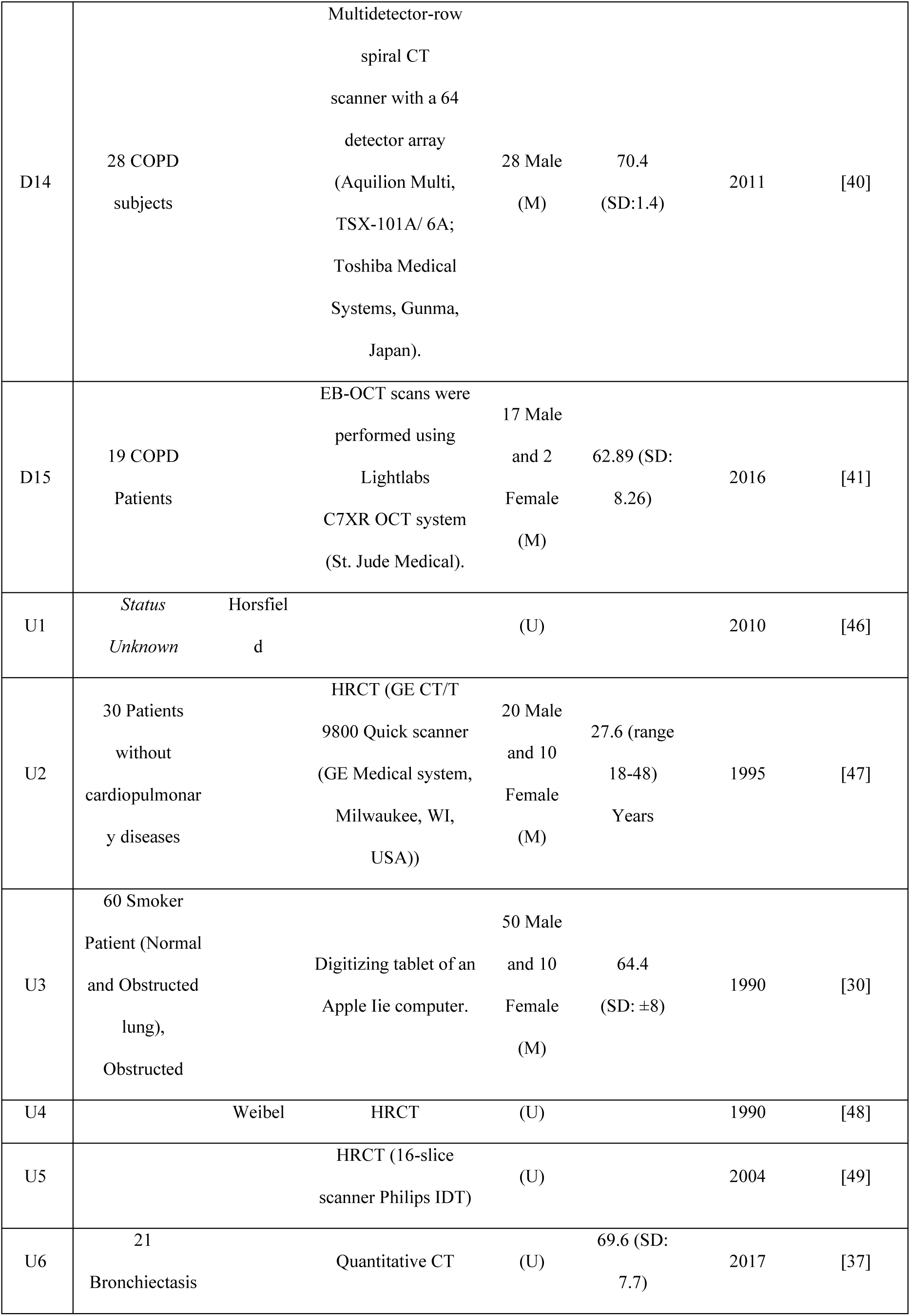
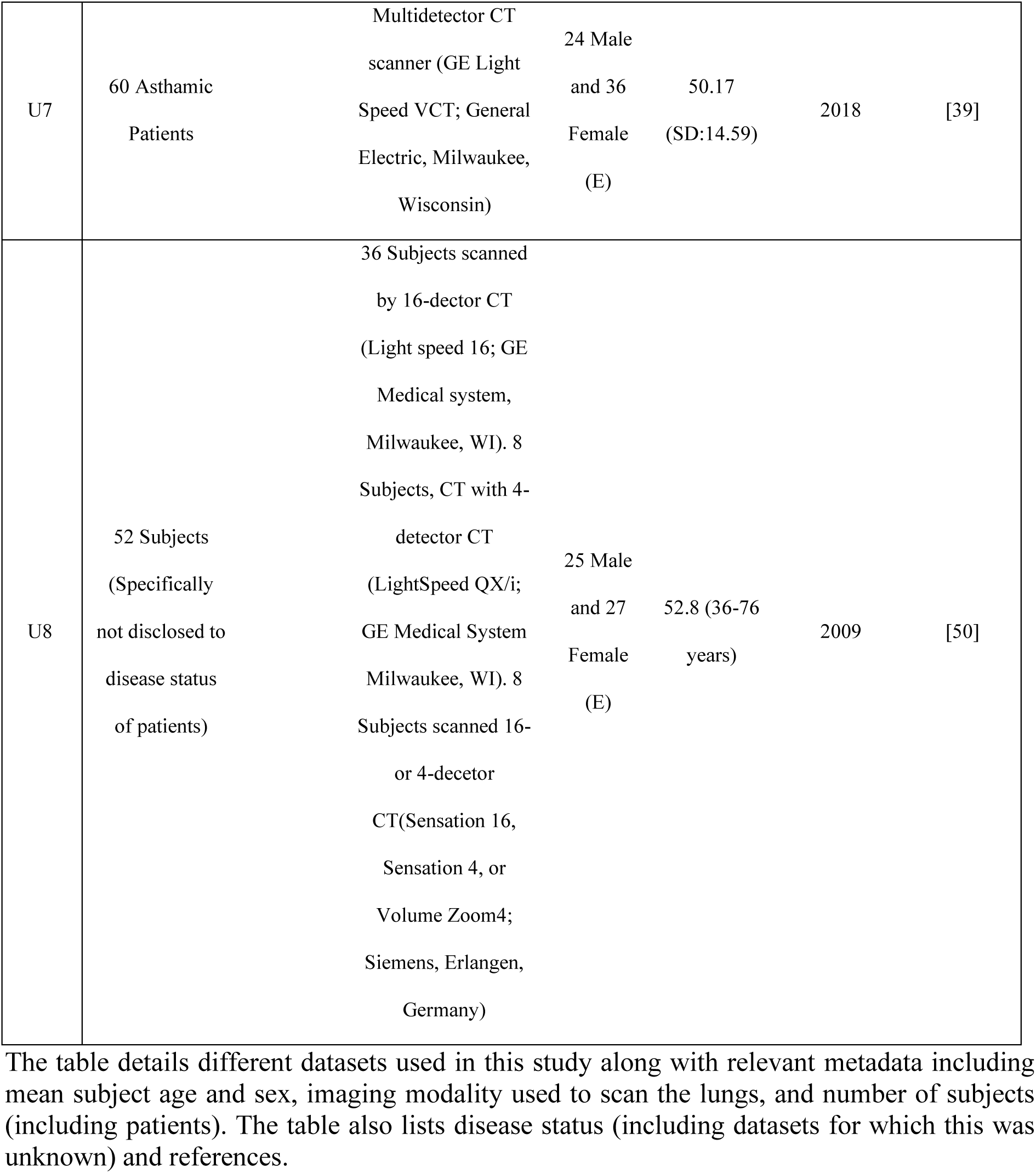
Summary of datasets, including their metadata, reviewed in this study.

We first investigated the mean age of the subjects/patients to explain the variance. We noted that while there was a general trend in both health and disease that the wall thickness reduced with age (Figures 2A,B), the CoV was high for all categories except for the age category 20-35 in disease dataset for which only one study was available (this meant variance could not be calculated). The anatomically-relevant health/disease clusters showed no enrichment for a specific age category (Figure 2C,D). We also found that sex differences could not account for the lack of consistency, with both healthy and patient datasets showing no enrichment (Figure 2E,F). Though it must be added that we could only find two studies that were “female-dominated”. The date of publication yielded similar trends as well.

**Figure 2:**
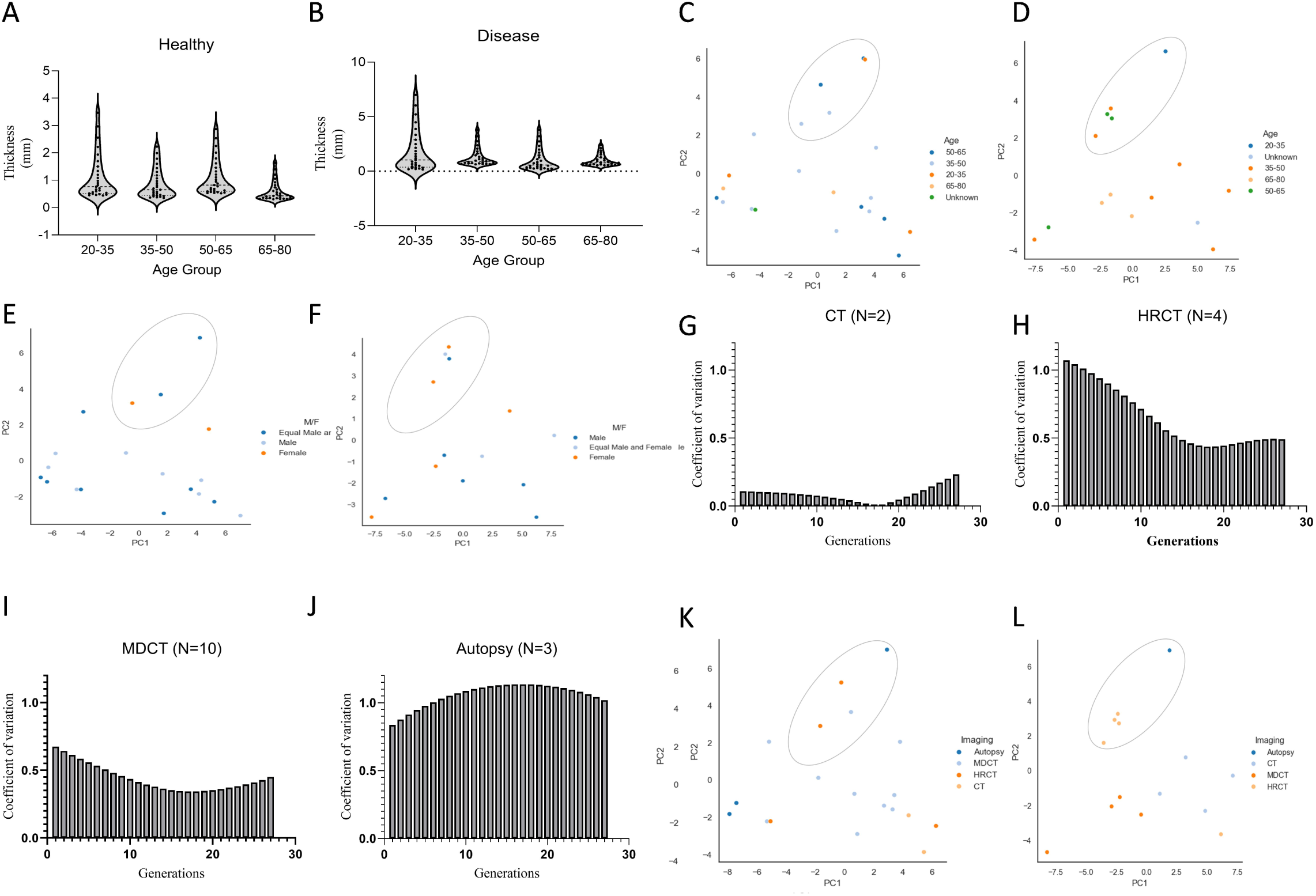
**(A–B)** Violin plot shows estimated airway wall thickness across various age groups for healthy (A) and diseased (B) datasets. Each dot represents an airway generation. **(C–F)** PCA projection with K-means clustering of wall thickness estimates for all 27 airway generations shown by age (C, D) and sex (E, F) for healthy (C, E) and diseased (D, F) datasets. **(G–J)** Histograms show the CoV for wall thickness estimates across all Horsfield airway generations for the healthy datasets based on the imaging modality: CT (G), HRCT (H), MDCT (I), and autopsy (J). **(K-L)** PCA projection with K-means clustering of wall thickness estimates for all 27 airway generations shown by imaging modality for healthy (K) and diseased (L) datasets.

We next investigated if the modality used for imaging subject/patient lungs can explain this variance. To achieve this, we first plotted the CoV for estimates generated via CT, HRCT, MDCT, autopsies (Figures 2G-J). The CoV was again high for all modalities, except CT. However, the studies that utilised CT were underpowered (n=2 datasets). We next considered the anatomically-relevant health/disease clusters. While the health clusters did not show enrichment for specific modality (Figure 2K), the anatomically disease cluster showed predominance of HRCT data points (Figure 2L), suggesting the imaging modality can potentially explain variance. To explain the lack of result in the health cluster, we conducted the PCA again but on thickness estimates on airway generations >2 mm in diameter, and assessed the cluster at PC1 ∼0, and PC2 >2 (given these PC coordinates had yielded anatomically-relevant clusters initially in Figures 1F,G). This cluster did not show enrichment for a specific modality. To ensure consistency, we assessed whether this cluster showed a higher proportion of for other attributes (sex, age, date of publication attributes), which was not the case.

### Inconsistency in wall thickness estimates significantly impacts lung function predictions

We next quantified what, if any, impact the inconsistency in reported wall thickness estimates has on lung function predictions. We achieved this by selecting estimates representing each of the three health clusters, using these estimates to inform the development of virtual lung geometries, and simulating an extended (see Methods) Swan *et al*. (2012) [7] ventilation model to quantify the impact on lung function. Specifically, we used the following studies reporting healthy subjects; *Cluster I*: H3 and H10; *Cluster II*: H6 and H13; and *Cluster III*: H1 and H17. The model calculated airway resistance in parallel throughout the entire airway tree, producing significantly different lung function outcomes across the three Clusters. Airway resistance calculations vary from 0.448 cmH20/L/s to 2.133 cmH20/L/s, which represents >375% difference from the original [7] ventilation model output value. Next, we plotted the pressure-volume curves (Figure 3A).

**Figure 3:**
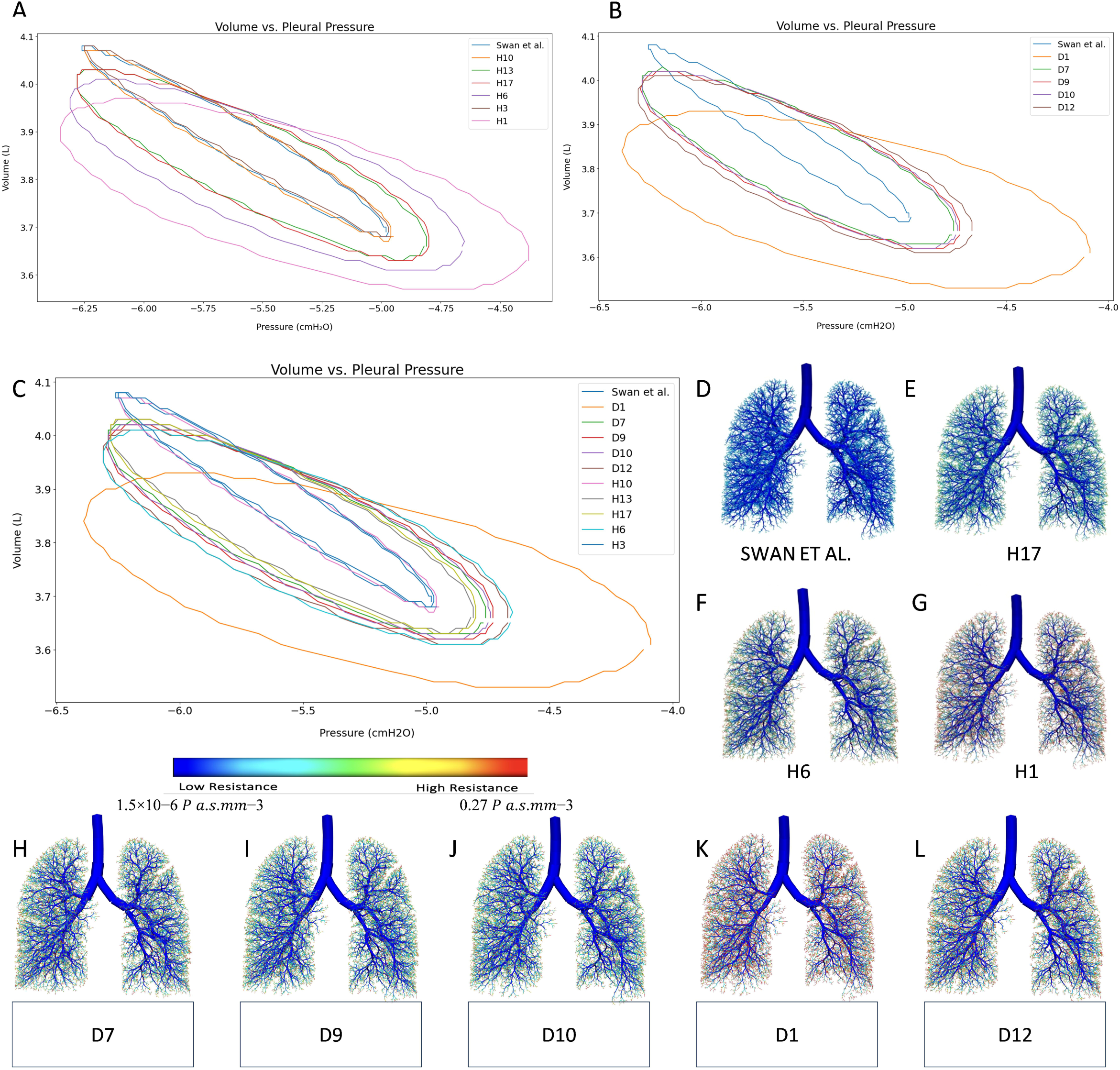
**(A–C)** Volume vs pleural pressure curves for selected healthy (A) and diseased (B) datasets, which have been combined in (C). **(D–L)** Airway resistance predicted for respective studies based on the estimated dimensions and compared against Swan *et al*. (2012). Colour scale indicates airway resistance in Pa.s/mm^3^.

Pressure-volume curve shape, size, and tilt angle are indicative of several aspects of lung function, such as total lung compliance and work of breathing. Figure 3A shows that the curves split into four groups. The first group is formed of H10 and H3 (i.e. *Cluster I* studies) along with the original lung ventilation model [7] outputs, the second by H13, H6, H1 and H17 (*Cluster II*). The groups were observed to approximately share peak volume, degree of hysteresis, compliance and work of breathing. Similarly, we simulated airway resistance and PV-curves for the studies reporting patient airway estimates. Specifically, we used the following studies; *Cluster II:* D7, D9, D10, D1 and D12. The pressure volume curves form three clusters; the first one including Swan *et al*. 2012 [7] reflecting healthy lung dimensions for relative comparisons. The second includes D7, D9, D10, and D12 and the third includes D1 (Figure 3B). The relative airway resistance was plotted in Figure 4H-L.

## DISCUSSION

Accurately estimating airway dimensions, especially at the patient-specific level, is critical to developing and deploying precision medicine strategies that aim to tailor therapies to patients’ unique profiles by first predicting the impact of multiple therapies on the lung function of individual patients [18]. Here, we asked if reported estimates of airway dimensions were consistent and whether simulations underpinned by these estimates to predict changes in lung mechanics and function will be clinically relevant? Our analysis revealed that reported airway dimensions lack consistency. PCA and unsupervised clustering revealed the data to group into multiple clusters (three for healthy subjects, four for diseased). In both analyses, we found that datasets that clustered around PC1 ∼0, and PC2 >2 yielded physiologically-relevant estimates. We investigated the source of this variance in both healthy and diseased datasets by conducting unsupervised clustering and assessing the enrichment of various clusters by attributes such as study subject age and sex, date of publication of study, and imaging modality used in the study. No health clusters were enriched for any factors, negating their contribution to the observed variance However, the physiologically-relevant cluster for the diseased dataset showed predominance of HRCT data points, weakly implicating the imaging modality for the inconsistency.

To quantify the impact of this inconsistency on lung function predictions, we used the wall thickness estimates from the various clusters to predict airway resistance and volume vs pleural pressure of lung airways. The simulations revealed that the resistance produced across the three ‘health’ clusters varied by >375% with downstream impact on hysteresis, compliance, and work of breathing. While wall thickness does not affect total lung compliance directly, the increased obstruction (given the choice of wall thickness estimates) prompts the model to settle at a lower peak volume. For clarity, the model was run for each wall thickness with the same tidal volume requirement to isolate the effect of wall thickness. Therefore, a lower peak volume causes a proportional decrease in the model’s estimate of FRC and acinar compliance. This observation emphasises that model functionally can be sensitive to wall thickness estimates, potentially affecting multiple lung function indices.

Clearly, the choice of airway dimension estimates heavily influences predictions generated from lung mechanics and function models. Yet, the clinical and computational community lacks guiding principles across clinically/physiologically-relevant airway dimension estimate generation and usage. To overcome this challenge, we developed The Human Airway Registry: an online database that can be used by respiratory clinicians and researchers to share imaging data and airway dimension estimates. Following open-access principles, users can register for a free account, and use the database to download available data and/or upload their own (in *csv* or *xlsx* format). The database dynamically conducts PCA/K-means clustering and G1 vs G27 analyses on all data available, including the one uploaded by the user. This helps the user identify whether their choice of airway dimensions are physiologically/anatomically relevant. Computational users can also download the anatomically relevant data to inform the development of virtual lung geometries to simulate their lung ventilation and mechanics models. Enabling simulation of airway resistance and volume-pressure curves automatically based on estimates uploaded on the portal will form part of the sequel.

We acknowledge limitations to this analysis. First, some datasets reported dimensions in a graphical format. To derive quantitative estimates, we used image recognition software to quantify dimensional values for each airway generation, which can potentially add some variability. Second, very few studies reported dimensions for all airway generations, which meant we had to inter-/extra-polate available data to generate standardised estimates. Third, we found significant differences in estimates across studies that used the same baseline images to estimate airway dimensions. For example, Kamm *et al*. (1999) [20] reported luminal diameter and wall area with thickness to bronchial radius. Their numbers were different from the source study [21]. Separately, Montaudon *et al*. (2009) [22] did not explicitly report units and we had to assume (based on careful analysis) that the reported dimensions were in centimetres. These inconsistencies can be a contributory factor behind the high variance we observed across the published datasets. We mitigated this challenge by first standardising all dimension estimates to the Horsfield order followed by unsupervised clustering, which revealed all anatomically-relevant datasets to cluster together. Despite the high inconsistency we were, thus, still able to identify datasets reporting clinically-applicable, physiologically-relevant airway dimension estimates.

## CONCLUSIONS

Reported estimates of airway dimensions show high inconsistency. Our analysis weakly implicates imaging modalities used to generate lung images for this variance. But, sub-optimal reporting of data makes it challenging to identify the key contributor(s). Overcoming this inconsistency requires standardised reporting of airway dimensions and quality assessments to ensure dimension estimates show physiological-relevance. For this, we make available a web database to enable the research community download existing datasets, share their (novel) datasets, and confirm their physiological relevance. We anticipate that offering easy access to physiologically-relevant airway dimensions and analytic pipeline via our database to enable standardised usage of airway dimensions for computational modelling will support the development of more accurate patient-specific virtual airway geometries and lung function predictions accelerating the development, validation, and deployment of precision medicine strategies to target asthma, specifically, and respiratory diseases, broadly.

## Authors Contributions

Conceptualisation: MKO and HK. Methodology, Analysis, Visualisation: All authors. Software: RAS, MKO, SKM, HK. Data Curation: MKO and HK. Supervision: CEB, KB, HK. Writing—original draft: MKO, RAS, SKM, MS, KB, HK. Writing—review & editing: All authors. Funding: KB and HK.

## Competing interests

We declare no competing interests.

## Funding and Acknowledgments

The authors acknowledge Institute of Precision Health at the University of Leicester for the Future 100 PhD scholarship (MKO), the Royal Society of New Zealand via a James Cook Fellowship (KB) and the Dines Charitable Trust (supporting MS), EPSRC-funded NetworkPlus “Integrating BIOphysical data-driven modelling into REspiratory MEdicine - BIOREME” Grant #EP/W000490/1 (HK), UKRI CRCRM Grant #MR/Z505638/1 (HK), Royal Academy of Engineering Grant #RF\201920\19\275 (HK).

## Notes

### Competing Interest Statement

The authors have declared no competing interest.

